# The euchromatic histone mark H3K36me3 preserves heterochromatin through sequestration of an acetyltransferase complex in fission yeast

**DOI:** 10.1101/738096

**Authors:** Paula R. Georgescu, Matías Capella, Sabine Fischer-Burkart, Sigurd Braun

**Affiliations:** Department of Physiological Chemistry, BioMedical Center (BMC), Ludwig Maximilians University of Munich, Martinsried, Germany

**Keywords:** chromatin, heterochromatin, silencing, acetyltransferase, histone modification

## Abstract

Maintaining the identity of chromatin states requires mechanisms that ensure their structural integrity through the concerted actions of histone modifiers, readers, and erasers. Histone H3K9me and H3K27me are hallmarks of repressed heterochromatin, whereas H3K4me and H3K36me are associated with actively transcribed euchromatin. Paradoxically, several studies have reported that loss of Set2, the methyltransferase responsible for H3K36me, causes de-repression of heterochromatin. Here we show that unconstrained activity of the acetyltransferase complex Mst2C, which antagonizes heterochromatin, is the main cause of the silencing defects observed in Set2-deficient cells. As previously shown, Mst2C is sequestered to actively transcribed chromatin via binding to H3K36me3 that is recognized by the PWWP domain protein Pdp3. We demonstrate that combining deletions of *set2*^+^ and *pdp3*^+^ results in an epistatic silencing phenotype. In contrast, deleting *mst2*^+^, or other members of Mst2C, fully restores silencing in Set2-deficient cells. Suppression of the silencing defect in *set2*Δ cells is specific for pericentromeres and subtelomeres, which are marked by H3K9me, but not seen for loci that lack genuine heterochromatin. Although Mst2 catalyzes acetylation of H3K14, this modification is likely not involved in the Set2-dependent pathway due to redundancy with the HAT Gcn5. Moreover, while Mst2 is required for acetylation of the H2B ubiquitin ligase Brl1 in euchromatin, we find that its role in heterochromatin silencing is not affected by Brl1 acetylation. We propose that it targets another, unknown substrate critical for heterochromatin silencing. Our findings demonstrate that maintenance of chromatin states requires spatial constraint of opposing chromatin activities.

---

The stable maintenance of functional chromatin states is crucial for cellular identity. The nucleus of eukaryotic cells is organized into topologically distinct chromatin domains, known as eu- and heterochromatin, which are associated with different nuclear functions. Both types of chromatin are controlled through various post-translational histone modifications, nucleosome remodeling and RNA-related processes. Euchromatin is associated with active transcription and hyperacetylated, contributing to an open chromatin structure. In contrast, heterochromatin is associated with gene repression and hypoacetylated, often adopting a compact chromatin structure that restricts transcription and prevents genomic rearrangements. Constitutive heterochromatin is present at repeat-rich sequences that are part of specific chromosomal domains, like centromeres and telomeres, and plays a crucial role in genome integrity and stability. However, so-called facultative heterochromatin can also form at gene-rich regions, e.g. during cellular differentiation and adaptation to environmental changes. Responding to these changes and maintaining the structural integrity of heterochromatin domains once established requires the concerted actions of histone modifiers, readers, and erasers (Cavalli & Heard, 2019; Allshire & Madhani, 2018; Grewal & Jia, 2007).

A conserved type of heterochromatin is characterized by the presence of methylated Lys-9 of histone H3 (H3K9me) (Allshire & Madhani, 2018). This repressive mark is recognized by members of the Heterochromatin Protein 1 (HP1) family, which bind to di- and trimethylated H3K9 (H3K9me2/3) via their N-terminal chromodomain. HP1 molecules can dimerize through their C-terminal chromo shadow domains, which also participate in the recruitment of the H3K9 methyltransferase. This has been suggested to create a positive feedback loop, resulting in extensive spreading of H3K9me/HP1 and the establishment of a recruitment platform for other repressive factors. These include histone deacetylases (HDACs), which restrict histone turnover and chromatin accessibility through deacetylation of lysine residues at N-terminal histone tails, thereby contributing to the repressive state of heterochromatin. Those and other chromatin regulating pathways are conserved in the fission yeast *Schizosaccharomyces pombe* (*S. pombe*), which makes it a powerful model system to study the molecular mechanisms of heterochromatin establishment and maintenance (Allshire & Ekwall, 2015). Distinct heterochromatin domains are present at the sub-telomeres downstream of the telomeric repeats, the silent mating type locus, and the pericentromeric *dg* and *dh* sequences of the outer repeats (*otr*). H3K9me is deposited by a sole histone methyl-transferase, Clr4, which is present in a complex known as CLRC and catalyzes all three steps of methylation (mono-, di- and trimethylation) (Nakayama *et al*, 2001; Allshire & Ekwall, 2015; Iglesias *et al*, 2018). The repressive H3K9me mark is recognized by chromodomain-containing proteins, such as Clr4 itself (Zhang *et al*, 2008) and the HP1 homologs Swi6 and Chp2 that interact with various chromatin factors including the repressor complex SHREC (Motamedi *et al*, 2008; Fischer *et al*, 2009). This complex comprises a Snf2-like nucleosome remodeler (Mit1) and an HDAC (Clr3) subunit, analogous to the mammalian NuRD complex, and restricts access of RNA polymerase II (RNAP II) to heterochromatin (Sugiyama *et al*, 2007; Job *et al*, 2016; Leopold *et al*, 2019). Clr3 specifically deacetylates histone H3 at lysine K14 (H3K14ac), whereas other HDACs, such as Sir2 and Clr6, show a preference for H3K9ac (Wiren *et al*, 2005). Similar to Clr3, Clr6 is part of a multimeric complex and associates with Swi6 (Fischer *et al*, 2009). Both Clr3 and Clr6 are not restricted to heterochromatin but have global functions in histone deacetylation (Nicolas *et al*, 2007; Wiren *et al*, 2005).

Heterochromatin assembly is guided by several targeting mechanisms among which the RNA interference machinery plays a prominent role in *S. pombe* (Verdel *et al*, 2004; Martienssen & Moazed, 2015). Small interfering RNAs (siRNAs) produced from heterochromatic DNA repeats are loaded onto the Argonaute-containing RNA-induced transcriptional silencing (RITS) complex, which targets complementary nascent transcripts (Verdel *et al*, 2004). RITS further interacts with the RNA-dependent RNA polymerase complex (RDRC), leading to the generation of dsRNAs that are processed by Dicer into siRNAs and fed into the amplification loop (Motamedi *et al*, 2004). In addition, RITS recruits CLRC via the bridging factor Stc1 (Bayne *et al*, 2010), resulting in H3K9me deposition. Conversely, stable binding of RITS to heterochromatin is enhanced through recognition of H3K9me by the chromodomain protein Chp1, which is part of RITS (Sadaie *et al*, 2004; Schalch *et al*, 2009; Verdel *et al*, 2004). The physical coupling of siRNA processing and histone modification generates a reinforcing feedback loop that persists over multiple generations, allowing the stable epigenetic inheritance of heterochromatin domains (Yu *et al*, 2018; Kowalik *et al*, 2015; Duempelmann *et al*, 2019).

Conversely, euchromatin is protected from ectopic heterochromatin assembly by several heterochromatin-antagonizing factors. The JmjC protein Epe1 counteracts H3K9me formation at euchromatic sites prone to assembling heterochromatin (Ragunathan *et al*, 2015; Audergon *et al*, 2015; Zofall *et al*, 2012). Furthermore, it prevents heterochromatin spreading beyond its natural boundaries (Ayoub *et al*, 2003). Epe1 is recruited to HP1 proteins and competes with SHREC for HP1 binding, thereby facilitating access of RNAP II to chromatin (Zofall & Grewal, 2006; Shimada *et al*, 2009; Barrales *et al*, 2016). Conversely, the association of Epe1 with heterochromatin is restricted on one hand through HP1 phosphorylation, which favors SHREC recruitment (Shimada *et al*, 2009), and on the other hand through ubiquitin-dependent degradation, which confines Epe1 to the heterochromatin boundaries (Braun *et al*, 2011). Heterochromatin is further antagonized by the RNA polymerase II-associated factor 1 complex (Paf1C), which is involved in multiple steps in transcription. Mutants of Paf1C are susceptible to siRNA-mediated stochastic heterochromatin initiation at ectopic sites, possibly due to altered kinetics in the processing and termination of nascent transcripts (Kowalik *et al*, 2015; Yu *et al*, 2018; Shimada *et al*, 2016). Besides initiation, Paf1C affects heterochromatin maintenance. The Paf1C subunit Leo1 prevents spreading at heterochromatin boundaries and promotes histone turnover (Verrier *et al*, 2015; Sadeghi *et al*, 2015). Paf1C also seems to be important to overcome the repressive activity of H3K9me3, which has been linked to its elongation promoting function by which it may help RNAP II to more easily disrupt nucleosomes (Duempelmann *et al*, 2019).

Histone acetyltransferases (HATs) also counteract heterochromatin, either directly by altering the nucleosomes’ charge and structure, or indirectly through the recruitment of factors to acetylated histones. HATs can similarly modify the structure and activity of non-histone targets. The HAT Mst2 mediates acetylation of H3K14 redundantly with the SAGA complex subunit Gcn5 (Wang *et al*, 2012). Loss of Mst2 enhances silencing at subtelomeres (Gómez *et al*, 2005) and bypasses the need for RNAi in centromeric heterochromatin maintenance (Reddy *et al*, 2011). The absence of Mst2 aggravates the phenotype of Epe1-deficient cells, resulting in severe growth defects due to ectopic silencing of essential genes via H3K9me spreading (Wang *et al*, 2015). Furthermore, the rate at which ectopic silencing is initiated in a *paf1* mutants is drastically increased when the HAT Mst2 is absent (Flury *et al*, 2017). Mst2 is present in a complex (Mst2C) homologous to *S. cerevisiae* NuA3b, which contains the PWWP domain protein Pdp3 (Wang *et al*, 2012; Gilbert *et al*, 2014). Pdp3 binds to trimethylated H3K36 (H3K36me3) and sequesters Mst2 to actively transcribed chromatin (Gilbert *et al*, 2014; Flury *et al*, 2017). Notably, in Pdp3-deficient cells, Mst2 is no longer stably bound to transcribed genes but gains promiscuous access to other chromatin regions including heterochromatin, where it triggers a silencing defect (Flury *et al*, 2017). However, none of these heterochromatin-associated phenotypes are recapitulated by the loss of Gcn5, implying that Mst2 has another, non-redundant function that involves an acetylation substrate other than H3K14 (Flury *et al*, 2017; Reddy *et al*, 2011). Indeed, proteome analysis revealed that Mst2 is involved in the acetylation of Brl1, which is part of the histone ubiquitin E3 ligase complex (HULC). Replacing *brl1*^+^ with an acetylation-deficient mutant (*brl1-K242R*) recapitulated the increased rate of ectopic heterochromatin assembly when combined with a mutant of Paf1C, whereas employing a mutant that mimics acetylation (*brl1-K242Q*) completely restored the wild-type phenotype (Flury *et al*, 2017). However, whether Brl1 acetylation is also responsible for the silencing defect under conditions when Mst2 encroaches on heterochromatin (i.e. in *pdp3*Δ cells) remains unknown.

H3K36 methylation is associated with actively transcribed chromatin. In budding and fission yeast, all three methylation states are mediated by a single enzyme, Set2 (Wagner & Carpenter, 2012). Set2 binds to the phosphorylated C-terminal domain (CTD) of transcribing RNAP II through its Set2 Rpb1 interacting (SRI) domain, which is a prerequisite for H3K36 tribut not dimethylation (Suzuki *et al*, 2016; Tanny, 2014). H3K36me is recognized by various reader proteins through chromodomains, PHD fingers, or PWWP domains. While H3K36 methylation is coupled to transcriptional elongation, it is also implicated in gene repression. In budding yeast, the HDAC complex Rpd3S binds to chromatin via its chromodomain subunit Eaf3, which recognizes di- and trimethylated H3K36. Rdp3S recruitment and histone deacetylation in the wake of transcribing RNAP II prevents initiation of aberrant transcription from cryptic promoters within coding regions (Carrozza *et al*, 2005; Keogh *et al*, 2005; Joshi & Struhl, 2005). The fission yeast homologs of Rpd3S and Eaf3 are Clr6 complex II (Clr6C-II) and Alp13, respectively (Nicolas *et al*, 2007). Mutants deficient in Set2 and Alp13 display increased antisense transcription in coding regions (Nicolas *et al*, 2007) and silencing defects at various heterochromatin domains (Creamer *et al*, 2014; Suzuki *et al*, 2016; Chen *et al*, 2008). A requirement for Set2 was further reported for repressed subtelomeric regions that are characterized by highly condensed chromatin bodies termed ‘knobs’ that lack H3K9me and most other histone modifications (Matsuda *et al*, 2015). However, it remains unclear whether these silencing defects in *set2*Δ cells are mediated directly through a local loss of H3K36me, resulting in reduced binding of Clr6C-II to chromatin and increased histone acetylation, or through an alternative mechanism.

Here we demonstrate that the silencing defects in *set2*Δ cells can be fully reversed at repressed chromatin regions that are marked by canonical heterochromatin marks and are largely devoid of H3K36me3 by concomitant deletion of *mst2*^+^. Full suppression of the silencing defect is also seen for other Mst2C members that are critical for proper complex assembly. However, suppression is not observed for the PWWP subunit Pdp3 that recruits Mst2C to actively transcribed regions via H3K36me3. This strongly suggests that the silencing defect of *set2*Δ cells is caused indirectly by global mislocalization of Mst2C and its encroachment on heterochromatic regions. Conversely, other loci upregulated upon the loss of Set2 are not suppressed by *mst2*^+^ deletion, indicating that their repression is mediated by a different mechanism.

## Results

### Deletion of the mst2^+^ gene suppresses the silencing defect of Set2-decifienct cells

We previously showed that loss of the PWWP subunit Pdp3 causes moderate silencing defects for the pericentromeric *imr1L::ura4*^+^ reporter gene and various subtelomeric genes. These silencing defects are suppressed when *mst2*^+^ is concomitantly deleted (Flury *et al*, 2017). Since Pdp3 anchors Mst2 to euchromatin via H3K36me3 (Flury *et al*, 2017), we tested whether the silencing defects observed at various genomic loci in *set2*Δ cells (Matsuda *et al*, 2015; Creamer *et al*, 2014; Suzuki *et al*, 2016; Nicolas *et al*, 2007; Chen *et al*, 2008) can be attributed to Mst2C mislocalization. This hypothesis makes the prediction that silencing will be restored when Mst2 is eliminated in *set2*Δ cells, analogous to *mst2*^+^ deletion in a *pdp3*Δ strain (see scheme, Figure 1A). Silencing can be monitored *in vivo* using reporter assays with the *ura4*^+^ gene inserted into a heterochromatic region. Presence of the nucleotide analog 5-FOA (fluoroorotic acid) inhibits cell growth due to the conversion of 5-FOA into a toxic metabolite by the gene product of *ura4*^+^ but allows growth when *ura4*^+^ transcription is repressed. By examining pericentromeric silencing in the *imr1L::ura4*^+^ reporter strain used previously (Flury *et al*, 2017), we found that, similar to *pdp3*Δ cells and consistent with other studies that have reported silencing defects for *set2*Δ cells, growth of *set2*Δ cells on 5-FOA is impaired (Nicolas *et al*, 2007; Chen *et al*, 2008; Suzuki *et al*, 2016; Creamer *et al*, 2014). Remarkably, while cell growth in the presence of 5-FOA was not affected by loss of Mst2, it was nearly restored when *mst2*^+^ was deleted in a *set2*Δ background (Figure 1B). We confirmed the findings of the reporter assay by reverse transcriptase assays combined with quantitative PCR (RT-qPCR). Deletion of *set2*^+^ causes a reproducible upregulation of the *imr1L::ura4*^+^ reporter gene (4-fold) and two endogenous transcripts from the outer *dg* and *dh* repeats (both 3-fold; Figure 1C, left panels). In contrast, transcript levels in the *set2*Δ *mst2*Δ double mutant resemble those of wild-type (WT) cells. In addition, we examined expression levels of transcripts derived from a subtelomeric region that is marked with high levels of H3K9me2 (10-30 kb distal of the telomeric repeats). De-repression of the subtelomeric genes *tlh1*^+^*/tlh2*^+^, *SPAC212.09c* and *SPAC212.08c* in *set2*Δ cells is even more pronounced than what we found at peri-centromeres (10-, 15- and 60-fold, respectively; Figure 1C, right panels). Nonetheless, transcriptional upregulation at these loci is completely suppressed when *mst2*^+^ is concomitantly deleted in *set2*Δ cells.

**Figure 1:**
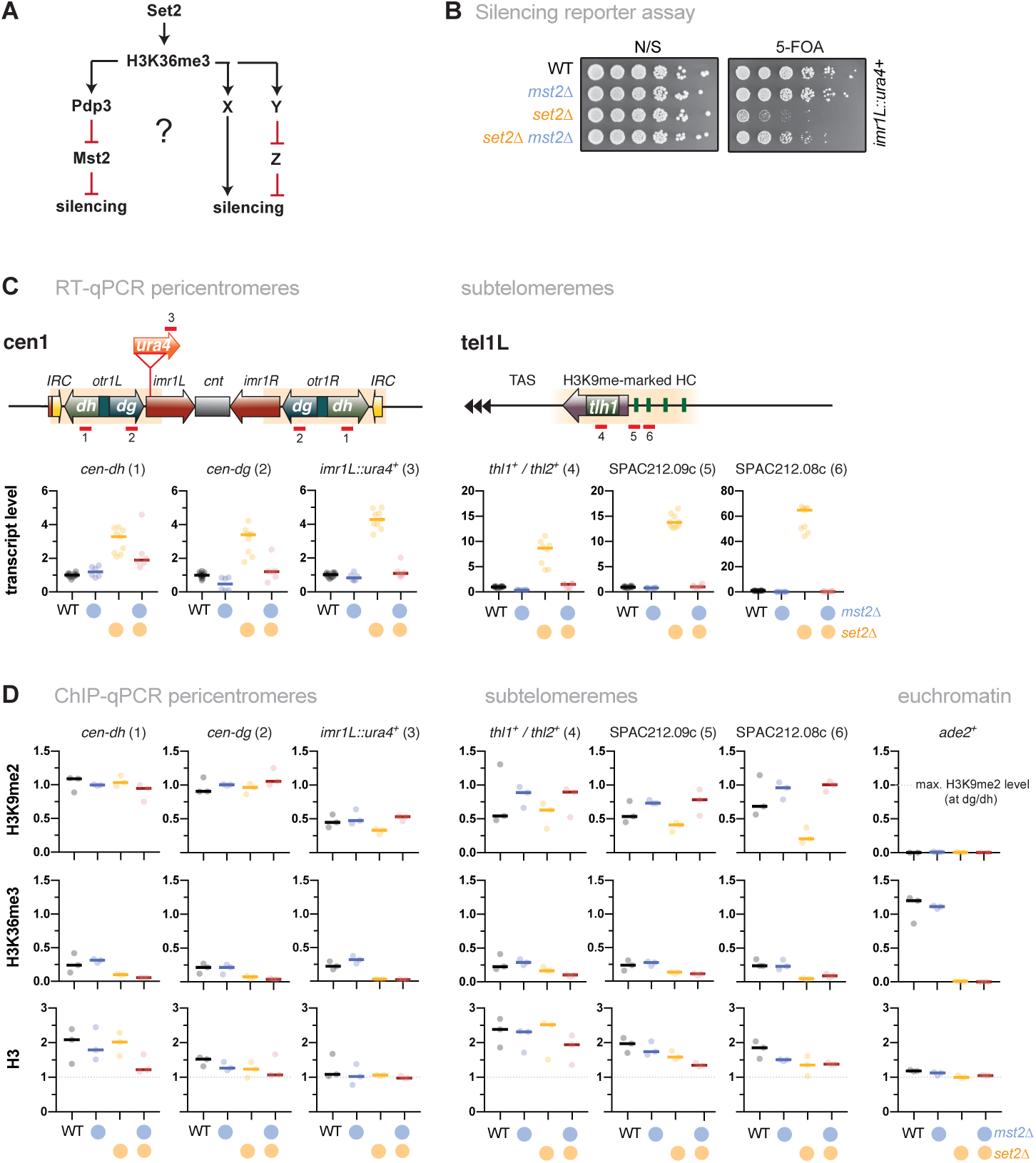
Loss of Mst2 recues the silencing defect caused by *set2*^+^ deletion. (A) Scheme depicting genetic interactions of *set2*^+^, *pdp3*^+^ and *mst2*^+^ contributing to heterochromatic silencing and potential parallel pathways in which H3K36me3 may be also involved. Black lines indicate positive regulations, red lines indicate negative regulations. (B) Silencing reporter assay with the *imr::ura4*^+^ reporter. Fivefold serial dilutions of wild-type (WT) cells and single and double deletion mutants of *mst2*^+^ and *set2*^+^; (N/S) nonselective medium. (C) RT-qPCR analysis. Shown are heterochromatic transcript levels of the strains used in (B). The schemes display the positions of the *ura4*^+^ reporter insertion and endogenous heterochromatic transcripts from pericentromeric (left) and subtelomeric heterochromatin (right); transcript levels have been normalized to *act1*^+^ and are shown relative to WT for each transcript. Circles and horizontal lines represent individual data and median from 6-12 independent experiments. (D) ChIP-qPCR analysis for H3K9me2 (top), H3K36me3 (middle) and H3 (bottom) at pericentromeric and subtelomeric heterochromatin; *ade2*^+^ (right panels) was used as control for euchromatin. Circles and horizontal lines represent individual data and median from 3 independent experiments. Input-normalized ChIP data were corrected for variation in IP efficiency by normalizing to the mean of *cen-dg* and *cen-dh* for H3K9me2, or the mean of three euchromatic loci (*tef3*^+^, *ade2*^+^, *act1*^+^) for H3K36me3 and H3. Note that H3K9me2 is largely unaltered at the *dh* repeats in *set2*Δ (Suzuki et al., 2016).

Constitutive heterochromatin in *S. pombe* is marked by high levels of H3K9me but largely devoid of euchromatic histone modifications (i.e. H3K4me, H3K36me) (Chen *et al*, 2008; Cam *et al*, 2005). However, residual levels of H3K36me3 have been detected at pericentromeres and subtelomeres, particularly in S phase during which peri-centromeric repeats are preferentially transcribed (Chen *et al*, 2008; Suzuki *et al*, 2016). By performing chromatin immunoprecipitation coupled to quantitative PCR (ChIP-qPCR), we found a varying degree of H3K9me2 decrease in *set2*Δ cells for several heterochromatic loci that display intermediate H3K9me2 levels (*imr1L::ura4*^+^ at pericentromeres; *SPAC212.09c* and *SPAC212.08c* at sub-telomeres; Figure 1D, upper panels). Conversely, *mst2*Δ and *set2*Δ *mst2*Δ cells showed elevated levels at those loci, suggesting that Mst2 counteracts H3K9me2 in a chromatin context-dependent manner. At most heterochromatic loci tested, H3K36me3 is low, reaching only 10-20% of the enrichment observed at euchromatin (Figure 1D, middle panels; note that the absolute level might even be lower, since the anti-H3K36me3 antibody used here shows limited cross-reactivity with H3K9me2; Supplementary Figure S1). We also analyzed nucleosome abundance by examining (total) histone H3. While H3 ChIP enrichments tend to be lower in the mutants for some loci, most changes were not significant and did not reflect transcriptional upregulation or changes in histone modifications (Figure 1D, lower panels). Together, these findings suggest that the silencing defects at pericentromeric and subtelomeric heterochromatin are not primarily caused by local changes of H3K36me3 within heterochromatin (which is already low in WT cells). Rather, our results imply that they are triggered by the uncontrolled activity of Mst2, which in the absence of Set2 is no longer tethered to euchromatin and gains promiscuous genome-wide access to chromatin, including the heterochromatic regions.

### Set2 acts in the same genetic pathway as other Mst2C members

We previously showed that the Mst2C subunit Pdp3 mediates Mst2 recruitment via its PWWP domain that binds to H3K36me3 (Figure 2A, left panel) (Flury *et al*, 2017). Since Set2 acts upstream of Pdp3 in Mst2 recruitment, we tested whether Set2 and Pdp3 also participate in the same pathway with respect to heterochromatin silencing. In agreement with our previous findings (Flury *et al*, 2017), we discovered that cells lacking Pdp3 displayed alleviated silencing at pericentromeres and subtelomeres, although the transcriptional increase was less pronounced in *pdp3*Δ compared to *set2*Δ (Figure 2B and C, left panels). Combining both deficiencies did not result in an additive increase; we rather observed a mild suppressive phenotype for *set2*Δ *pdp3*Δ when compared to the *set2*Δ single mutant. Although the nature of the partial suppression remains unclear (see discussion), the non-additive phenotype of the double mutant suggests that Pdp3 and Set2 act in the same pathway.

**Figure 2:**
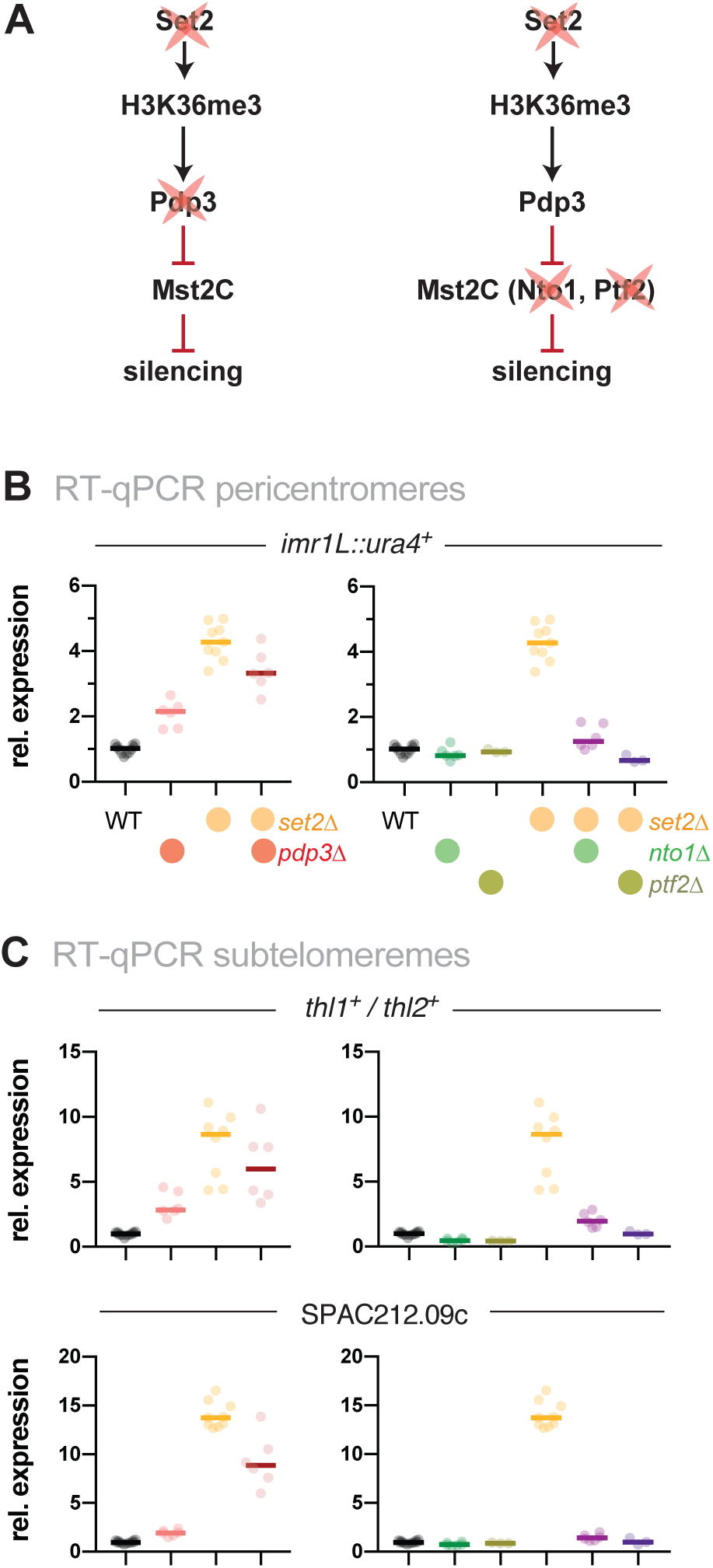
Loss of heterochromatin silencing in *set2*Δ is dependent on functional Mst2C. (A) Scheme displaying genetic interactions and genes mutated for experiments shown in (B) and (C). (B) + (C) RT-qPCR analysis of transcript levels at pericentromeric (B) and subtelomeric HC (C) RT-qPCR data analysis and primer positions as in Figure 1C. Circles and horizontal lines represent individual data and median from 6 independent experiments (except WT: *n* = 12; *ptf2*Δ and *ptf2*Δ *set2*Δ: *n* = 3)

Besides Mst2 and Pdp3, Mst2C contains five additional subunits (Nto1, Eaf6, Tfg3, Ptf1 and Ptf2). The functions of these subunits are not well understood, but Nto1 and Ptf2 are essential for the integrity and assembly of the complex, and mutants lacking either of these subunits phenocopy the loss of Mst2 (Wang *et al*, 2012). We therefore tested whether absence of Nto1 or Ptf2, analogous to *mst2*^+^ deletion, suppresses the silencing defect of *set2*Δ (Figure 2A, right panel). In contrast to *pdp3*Δ, single mutants of *nto1*Δ and *ptf2*Δ did not display elevated heterochromatic transcripts at pericentromeres and subtelomeres; instead, like *mst2*Δ, these deletions completely suppressed the silencing defect of *set2*Δ (Figure 2B and C, right panels). From this we conclude that an intact Mst2 complex is required to trigger the silencing defect at heterochromatin.

### Silencing defects in set2Δ at other repressed loci involve an Mst2-independent pathway

Loss of *set2*^+^ does not only affect chromatin regions with high levels of H3K9me2 but also other subtelomeric loci. These include the telomere-associated sequences (TAS) and a region about 50 kb downstream of the telomeric repeats, which is characterized by highly condensed chromatin bodies dubbed ‘knobs’ (Matsuda *et al*, 2015) (Figure 3A-B). Transcription of the non-coding RNA *TERRA* (telomeric repeat-containing non-coding RNA) from the TAS is repressed by heterochromatin and members of shelterin, the telomere-end protecting complex (Bah *et al*, 2012; Greenwood & Cooper, 2012). However, this subtelomeric region displays low nucleosome abundance and establishes only little H3K9me (van Emden *et al*, 2019). Similarly, subtelomeric ‘knob’ genes are decorated with a low amount of H3K9me2, and H3K36me3 is also reduced compared to euchromatin (Figure 3C-D).

**Figure 3:**
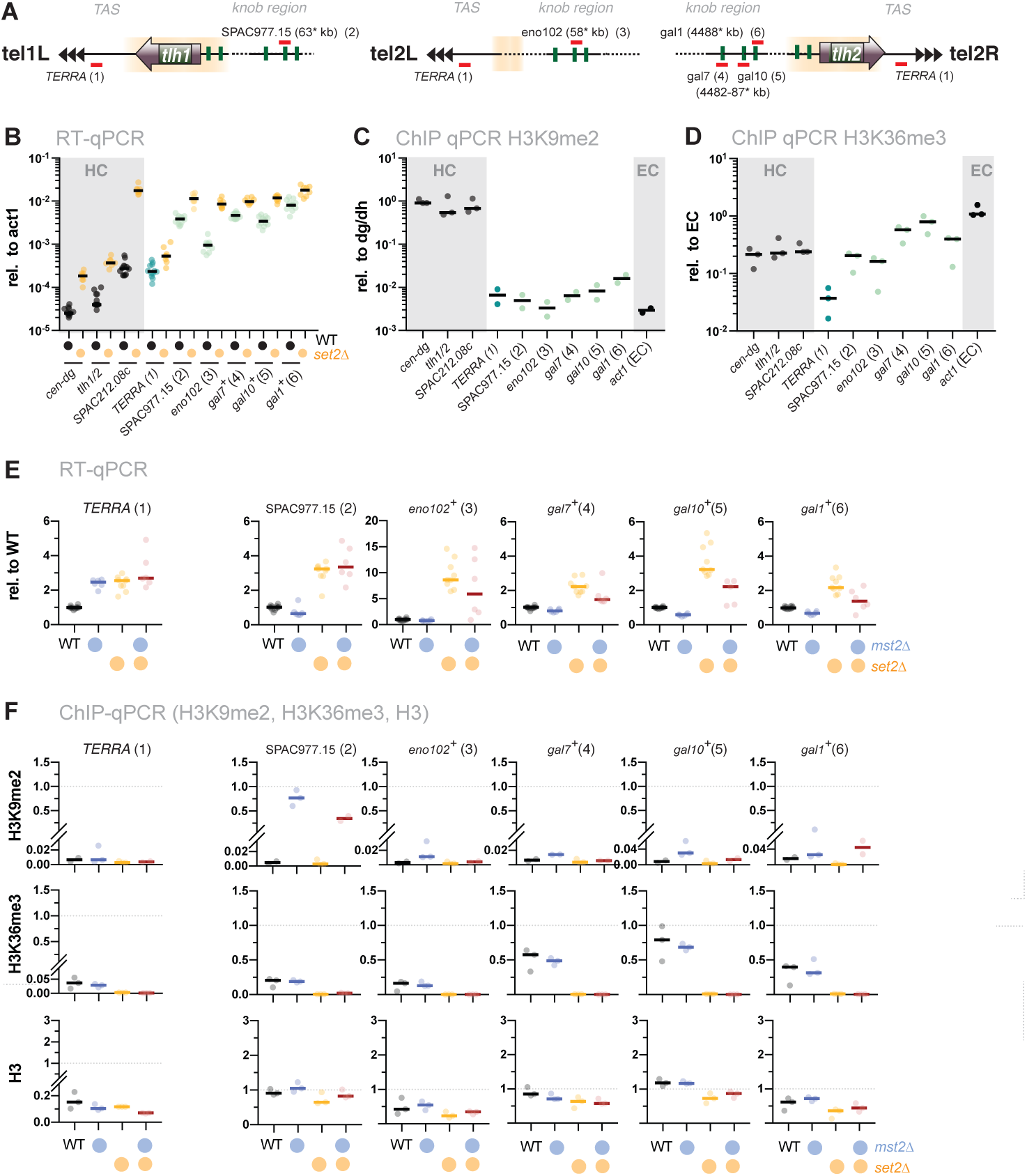
The silencing defect at ‘knobs’ in *set2*Δ is not associated with the Mst2 pathway. (A) Scheme depicting expression sites of the non-coding RNA *TERRA* and the positions of several loci within the ‘knob’ regions on chromosomes I and II. Chromosomal positions refer to annotations in www.pombase.org but differ from the absolute positions due to missing sequences at the chromosomal termini. (B) RT-qPCR analysis in WT and *set2*Δ strains comparing transcript levels at pericentromeric and subtelomeric HC (shaded in grey) to loci in the ‘knob’ region. Transcript levels from 12 independent experiments are shown relative to *act1*+. (C) + (D) ChIP-qPCR analysis of H3K9me2 and H3K36me enrichments in WT cells at loci analyzed in (B). Shown are ChIP data from 2-3 individual experiments analyzed as described in Figure 1D. (E) RT-qPCR analysis in *mst2*Δ and *set2*Δ single and double mutants; data from independent experiments analyzed as described in Figure 1C (WT *n* = 12; *mst2*Δ, *mst2*Δ *set2*Δ: *n* = 6; *set2*Δ: *n* =9). (F) ChIP-qPCR analysis of *TERRA* and different loci located within the ‘knob’ region; data from independent experiments analyzed as described in Figure 1D (*n* = 3).

When we examined expression of these sub-telomeric genes, we found a moderate but reproducible upregulation (3-fold) of *TERRA* in *set2*Δ. A similar increase was also observed in *mst2*Δ, and concomitant deletion of *mst2*^+^ in the *set2*Δ mutant did not suppress the silencing defect (Figure 3E, left panel). In agreement with previous studies, we also detected a 2 to10-fold upregulation for several ‘knob’ genes (Matsuda *et al*, 2015; Suzuki *et al*, 2016). As seen for *TERRA*, additional deletion of *mst2*^+^ did not restore silencing for these loci in *set2*Δ (or caused only a partial suppression; Figure 3E). Nonetheless, disruption of *mst2*^+^ promoted the establishment of H3K9me2 at several knob genes (particularly *SPAC977.15*), corroborating the notion of Mst2C playing a global role in antagonizing heterochromatin (Wang *et al*, 2015; Reddy *et al*, 2011; Flury *et al*, 2017). However, since removal of Mst2 was not sufficient to reinstate silencing in the absence of Set2, we presume that an additional, Mst2- and H3K9me-independent pathway, which likely does not rely on H3K9me, interferes with the repression of these genes.

### Mst2-dependent silencing defects are not mediated through Brl1 acetylation

We previously showed that Mst2 is involved in the acetylation of the non-histone substrate Brl1, a conserved ubiquitin E3 ligase that mono-ubiquitylates histone H2B at lysine 119 (Flury *et al*, 2017). Acetylation of Brl1 at lysine 242 (Brl1-K242ac) may have a stimulatory effect on its enzymatic activity (H2B-K119ub) and downstream events (H3K4me3), which protects euchromatic genes against the ectopic formation of heterochromatin, likely through increased transcription (Flury *et al*, 2017). We therefore wondered whether the Brl1-K242ac-dependent positive feedback loop is the main cause for the silencing defect observed in *set2*Δ cells. Analogous to suppression by the *mst2*^+^ deletion, we therefore combined the Set2 deficiency with the single-point mutant *brl1-K242R*, which mimics non-acetylated lysine (Figure 4A). Yet, in stark contrast to *set2*Δ *mst2*Δ cells (Figure 1C), we found that silencing at pericentromeres and sub-telomeres was not reinstated in the *set2*Δ *brl1-K242R* double mutant (Figure 4B-C). From this we conclude that preventing Brl1 acetylation is not sufficient to block the anti-silencing activity of Mst2 and that it targets at least one other substrate critical for heterochromatin silencing.

**Figure 4:**
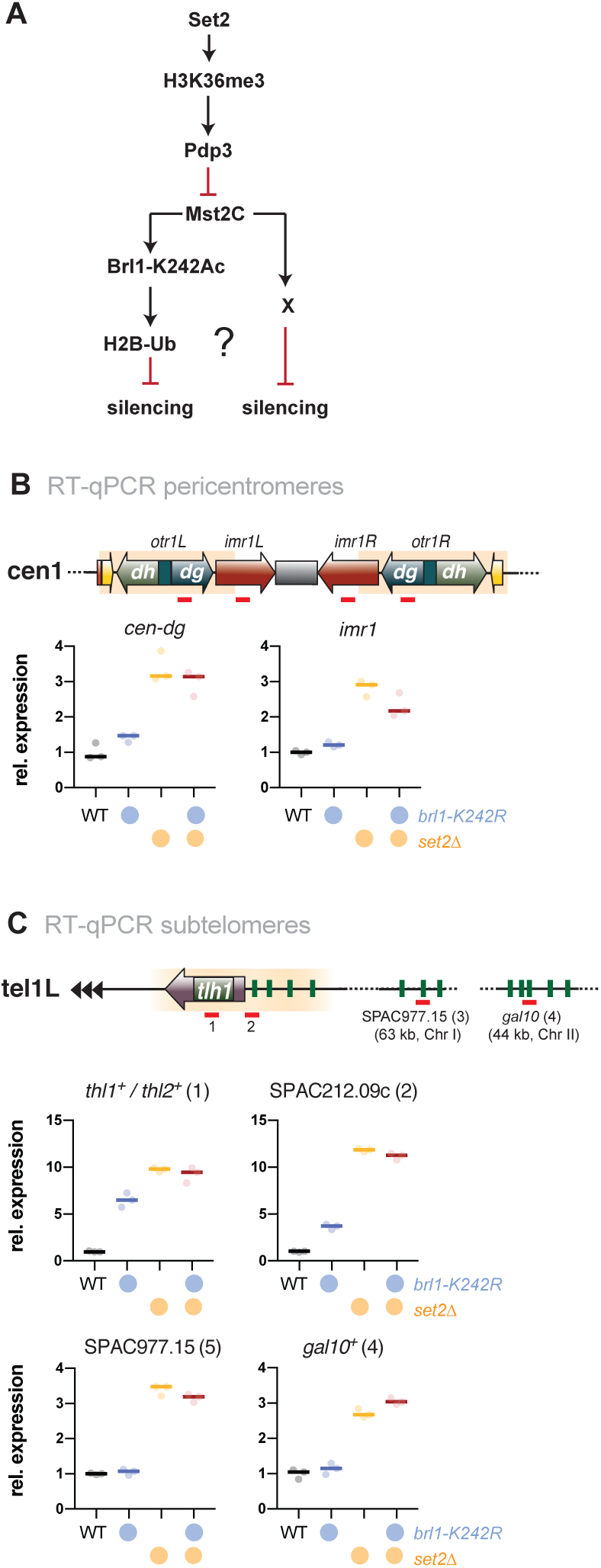
The target of Mst2 in heterochromatin is not Brl1. (A) Scheme depicting the described Mst2C pathway (Flury et al., 2017) involving Brl1-K242 acetylation and H2B ubiq-uitylation and a potential alternative pathway on HC silencing; black arrows represent positive regulation and red lines represent negative regulation. (B) + (C)) RT-qPCR analysis of transcript levels at pericentromeric HC (B) and subtelomeric HC and the ‘knob’ region (C). Data analysis as in Figure 1C (*n* = 3 individual experiments)..

## Discussion

### Set2-dependent silencing defects in heterochromatin are functionally linked to Mst2C

Transcriptionally active and repressed chromatin regions are marked by different posttranslational modifications. H3K36me3 in particular is deposited co-transcriptionally through the recruitment of Set2 by transcribing RNAPII and likely other elongating factors (Tanny, 2014). Yet paradoxically, loss of-Set2 also causes defects in transcriptionally repressed heterochromatin (Creamer *et al*, 2014; Suzuki *et al*, 2016; Chen *et al*, 2008; Nicolas *et al*, 2007). We previously have shown that deleting *mst2*^+^ restores silencing in cells lacking the Mst2C subunit and H3K36me3 reader Pdp3 (Flury *et al*, 2017). Here, we demonstrate that eliminating the catalytic HAT subunit Mst2, or other subunits critical for assembly of Mst2C, fully reverses the silencing defect of Set2-deficient cells at constitutive chromatin regions (Figure 1-2). Reinstatement of silencing is seen at the level of transcription as well as at the level of heterochromatin structure (H3K9me2) at those loci where lack of Set2 results in a partial decrease of both (Figure 1). In contrast, H3K36me3 is low at these heterochromatin regions in WT cells and not affected upon deletion of *mst2*^+^. This suggests that defective silencing is not directly caused by the loss of H3K36me within heterochromatin but rather indirectly through the promiscuous activity of Mst2C. The complete restoration of silencing further implies that Set2 exclusively controls silencing at constitutive heterochromatin through sequestration of Mst2 by H3K36me3. Importantly, while Mst2 is not detected by ChIP at transcribed chromatin when H3K36me3 is absent, it has still access to chromatin as demonstrated by DamID, resulting in transient binding and encroachment on heterochromatin (Flury *et al*, 2017).

Recruitment of Mst2 to euchromatin is mediated by the Mst2C subunit Pdp3, which binds to H3K36me3 via its PWWP domain. Consistently, lack of Pdp3, or a point mutation within its PWWP domain, also produces a defect in heterochromatin silencing (Flury *et al*, 2017). However, we noticed that the silencing defect in *pdp3*Δ is less pronounced than in *set2*Δ (Figure 2), despite the fact that silencing can be fully restored in the absence of Mst2 (see Figure 1). A possible explanation would be that Mst2 recruitment involves another H3K36me3-binding factor that acts redundantly with Pdp3. Indeed, the Mst2C subunit Nto1 contains two PHD fingers, and the *S. cerevisiae* homolog shows affinity for H3K36me3 (Shi *et al*, 2007). However, since Nto1 is essential for Mst2C assembly, a putative role in restricting Mst2 to euchromatin would be masked by the complete loss of HAT activity, thus resulting in deviating phenotypes for *pdp3*Δ and *nto1*Δ mutants. Alternatively, Pdp3 may also contribute to the stability or activity of the complex (at least in part), in addition to its function in H3K36me3 anchoring. This could explain the intermediate phenotype of *pdp3*Δ cells compared to *set2*Δ on one hand, and *mst2*Δ*/nto1*Δ on the other. More work will be needed to better understand the functions of the individual subunits of Mst2C.

### Set2-mediated gene repression at chromatin lacking H3K9me is independent of Mst2C

Besides defects at constitutive heterochromatin, deletion of *set2*^+^ results in the transcriptional up-regulation of other genes that are part of repressed chromatin regions but largely devoid of H3K9me2. These include the telomere-proximal gene encoding *TERRA* and various genes expressed from the subtelomeric ‘knob’ region, which is characterized by high chromatin condensation and the absence of most post-transcriptional histone modifications (Matsuda *et al*, 2015). However, except for at *TERRA*, we still detect some H3K36me3 at these chromatin regions (Figure 3). Transcriptional up-regulation of these loci is moderate in *set2*Δ and not suppressed by concomitant Mst2 elimination. Nonetheless, Mst2 may still gain access to these loci and antagonize heterochromatin formation, as seen by increased levels of H3K9me2 upon *mst2*^+^ deletion. However, the lack of suppression suggests the involvement of a Set2-reliant but Mst2-independent pathway that is critical for gene repression.

Another complex potentially involved in Set2-dependent silencing is the HDAC Clr6C-II, which contains the chromodomain protein Alp13 (Eaf3 in *S. cerevisiae*; MORF4 in humans) (Nicolas *et al*, 2007; Nakayama *et al*, 2003). The homologous complex in *S. cerevisiae*, Rpd3S, prevents cryptic transcription through deacetylation of histones in coding regions marked with H3K36me2/3 to which this complex is recruited via binding of Eaf3 (Carrozza *et al*, 2005; Keogh *et al*, 2005; Joshi & Struhl, 2005). Similarly, Clr6C-II promotes deacetylation of H3K9 in coding regions, and both Alp13 and Set2 prevent antisense transcription and repress peri-centromeric repeats (Nicolas *et al*, 2007). In addition, like H3K36me3, Alp13 accumulates on heterochromatin during S phase when pericentromeric repeats are transcribed (Chen *et al*, 2008). However, in contrast to its *S. cerevisiae* homolog, *S. pombe* Set2 does not contribute to deacetylation of bulk histones, only moderately affects antisense transcription, and displays an additive defect in a *set2*Δ *alp13*Δ double mutant, arguing for parallel pathways (Nicolas *et al*, 2007). Moreover, while H3K36me2 is sufficient to recruit budding yeast Rpd3S (Li *et al*, 2009), heterochromatin silencing and recruitment of Pdp3/Mst2C requires H3K36me3 (Flury *et al*, 2017; Suzuki *et al*, 2016). Thus, together with the fact that silencing is fully restored in the absence of Mst2, it appears less likely that Clr6C-II contributes significantly to the Set2-dependent pathway at heterochromatin. Still, it remains an attractive hypothesis that Clr6C-II represses transcription in an H3K36me/ Alp13-dependent manner at other loci where Mst2 plays a less prevailing role.

### Mst2C has distinct cellular functions by acetylating multiple targets

Mst2 acetylates H3K14 *in vitro* and *in vivo* and acts redundantly with the SAGA member Gcn5 (Wang *et al*, 2012). H3K14ac is critical for G2/M check-point activation upon DNA damage and controls chromatin compaction through recruitment of the nucleosome remodeler RSC (Wang *et al*, 2012). In addition, H3K14ac accumulates in heterochromatin upon deletion of the HDAC Clr3 and other components of the repressor complex SHREC, suggesting a function in antagonizing heterochromatin silencing (Grewal *et al*, 1998; Sugiyama *et al*, 2007). Furthermore, the anti-silencing factor Epe1 physically interacts with SAGA and targets the HAT to heterochromatin when Epe1 is overexpressed. This triggers a silencing defect that is accompanied by an H3K14ac increase and is dependent on the HAT activity of Gcn5 (Bao *et al*, 2018). At first glance, this seems reminiscent of the phenotype caused by relocalization of Mst2. However, additional findings cast doubt on whether H3K14 is the relevant substrate that mediates the anti-silencing function of Mst2. First, neither lack of Set2 nor of Pdp3 (both causing delocalization of Mst2) results in H3K14ac accumulation at heterochromatin (Flury *et al*, 2017; Suzuki *et al*, 2016). Second, while elimination of Mst2 bypasses the need for RNAi in pericentromeric silencing, this is not the case for mutants lacking Gcn5 (Reddy *et al*, 2011). Third, lack of Mst2, but not of Gcn5, promotes the assembly of ectopic heterochromatin domains (Flury *et al*, 2017). Together, these findings suggest a non-redundant function of Mst2 in antagonizing heterochromatin that likely involves another substrate than H3K14.

We previously showed that the euchromatin-protective role of Mst2 is mediated through acetylation of the non-histone substrate Brl1, a subunit of HULC (Flury *et al*, 2017). In particular, replacing *brl1*^+^ with an acetylation-deficient mutant (*brl1-K242R*) phenocopied the deletion of *mst2*^+^, whereas mimicking acetylation (*brl1-K242Q*) bypassed the need for Mst2 in protecting euchromatin. In agreement with this finding, deletion mutants of *brl1*^+^ or other components of HULC display more robust heterochromatin silencing than wild-type cells (Zofall & Grewal, 2007). Thus, acetylated Brl1 seemed to be a likely candidate for mediating the anti-silencing function of Mst2 also at heterochromatin. However, in contrast to *mst2*Δ, introducing the *Brl1-K424R* mutant was not sufficient to suppress the silencing defect of *set2*Δ. Instead, we observed transcriptional upregulation even in the single *brl1-K242R* mutant, suggesting that the loss of its acetylation target Brl1 (in euchromatin) renders Mst2 more active (in heterochromatin). While *set2*Δ cells may have various pleiotropic defects, we think that these are less likely the major cause of the silencing defect, since this phenotype can be fully reversed in the absence of Mst2. We therefore speculate that besides H3K14 and Brl1-K242 Mst2C targets at least one other substrate that is important for heterochromatin maintenance (Figure 5). Previous proteomics failed so far to identify additional acetylated substrates besides Brl1 that involve Mst2 (Flury *et al*, 2017). However, since heterochromatin makes up only a small portion of the genome and Mst2 may have only transient access to these genomic regions, a heterochromatin-specific substrate would be more difficult to identify.

**Figure 5:**
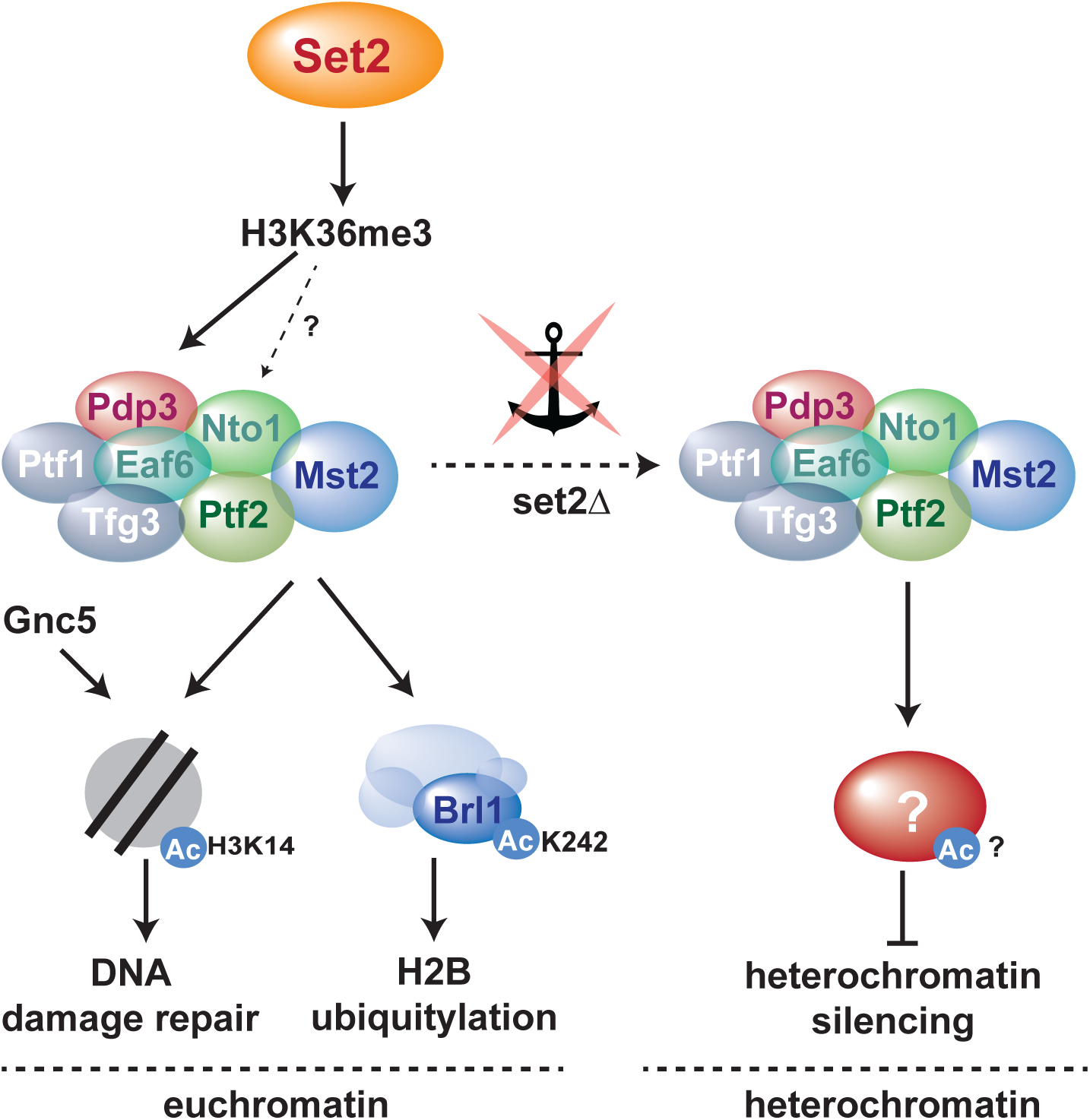
Model for Mst2C-dependent functional pathways. In the presence of H3K36me3, MstC is sequestered to euchromatin; in the absence of H3K36me3-mediated anchoring, Mst2C gets promiscuous access to heterochromatin. See text for details

### H3K36me-mediated HAT sequestration is conserved in heterochromatin maintenance

In worm embryos, perinuclear heterochromatin is established through two methyltransferases (MET-2, SET-25) and the nuclear membrane-associated chromodomain protein CEC-4, which tethers H3K9me-marked chromatin to the nuclear periphery (Gonzalez-Sandoval *et al*, 2015; Towbin *et al*, 2012). However, during differentiation, this pathway becomes redundant. An RNAi screen in *cec-4*-null larvae identified the chromodomain protein MRG-1 as a critical factor for perinuclear heterochromatin organization (Cabianca *et al*, 2019). MRG-1 is homologous to the Rpd3S subunit Eaf3 (*S. cerevisiae*) and Clr6C-II subunit Alp13 (*S. pombe*). In addition, MRG-1 also associates with the HAT CBP-1/p300. Although MRG-1 and CBP-1 are not homologous to Pdp3 and Mst2C, there are striking parallels (Cabianca *et al*, 2019): Like Pdp3, MRG-1 binds to H3K36me3-marked euchromatin. Loss of perinuclear heterochromatin in *mrg-1*-null is phenocopied by the double mutant lacking the H3K36 methyltransferases MET-1 and MES-4, whereas reducing CBP-1 restores silencing in *mrg-1-*null larvae. Overexpression of *cbp-1* is sufficient to release heterochromatin from the nuclear periphery in the absence of CEC-4. Moreover, the authors detected increased CBP-1 binding at several heterochromatic genes in *mrg-1*-null larvae. This was accompanied by elevated histone acetylation (H3K27ac), providing a direct link to gene expression.

Altogether, this demonstrates that the principle of heterochromatin maintenance through internal sequestration of HATs is conserved between fission yeast and worms, despite some apparent differences regarding the molecular mechanisms (i.e. the nature of the HAT enzymes and substrates). It further unveils that the pathways partitioning eu- and heterochromatin are remarkably entwined, requiring spatial constraint of opposing chromatin activities to maintain the identity of chromatin states. Recent observations have reinforced the notion that repressive histone marks contribute to epigenetic inheritance of chromatin domains (Ragunathan *et al*, 2015; Audergon *et al*, 2015; Yu *et al*, 2013; Duempelmann *et al*, 2019). In contrast, histone modifications associated with euchromatin have been considered rather a consequence than a cause of transcription. The discovery of H3K36me3 as a critical factor in heterochromatin maintenance will likely reopen the discussion to what extent ‘active’ marks also contribute to the epigenetic states of chromatin.

## Materials and Methods

### Contact for reagent and resource sharing

Important reagents and assays used are listed in Supplementary Table S5. Further information and requests for resources and reagents should be directed to Sigurd Braun.

### Yeast techniques and strains

Standard media and genome engineering methods were used (Fission Yeast. A Laboratory Manual. Hagan, Carr, Grallert and Nurse. Cold Spring Harbor Press. New York 2016). For the *ura4*^+^ reporter assay in Figure 1B cells were plated on EMM or EMM containing 1 mg/mL FOA. The strains were grown at 30°C for three (non-selective, NS) and four days (5-FOA), respectively. Cultures were grown at 30°C in liquid YES media (160 rpm, 12-24 hours) or at 30°C on solid YES agarose plates (for 3 days). The *brl1-K242R* point mutant was provided by M. Bühler (FMI, Basel). Strains used in this study are listed in supplementary Table S1

### RT-qPCR analyses

RT-qPCR experiments were carried out as previously described (Braun *et al*, 2011). The data are presented as individual data points together with the median. cDNA was quantified by qPCR using the primaQUANT CYBR Master Mix [Steinbrenner Laborsysteme] and a QuantStudio 5 Real-Time PCR System [Applied Biosystems] and primers listed in Supplementary Table S2. Prior to calculation of the median, *act1*^+^ normalized data sets from independent experiments were standardized to the mean of all samples from each experiment (experimental normalization; *eq 1.1*). These sample pool-normalized results were shown relative to the mean value of the sample pool-normalized wild type data from all (n) experiments (*eq 1.2*). Using the average from a collection (sample pool) instead of a single strain (e.g. WT) reduces bias, especially when transcripts levels are low in the repressed state and therefore more prone to noise.

### ChIP assays

ChIP experiments were performed essentially as described (Barrales et al. 2016). Cross-linking was performed with 1% formaldehyde for 10 min at RT. For quantitative ChIP, immunoprecipitations were performed with 2 µg of the following antibodies [cell lysates corresponding to different amounts of OD_600_]: anti-H3K9me2 [15 OD_600_] anti-H3K36me3 [5 OD_600_], and anti-H3 [5 OD_600_]. Antibodies are listed in Supplementary Table S4. Immunoprecipitated DNA was quantified by qPCR using the primaQUANT CYBR Master Mix [Steinbrenner Laborsysteme] and a QuantStudio 5 Real-Time PCR System [Applied Biosystems]. Primers are listed in Supplementary Table S2. Unless otherwise noted, the median was calculated from three independent experiments. qPCR signals were normalized against the input samples for each primer position as internal control. For ChIP experiments with anti-H3K9me2, the input-normalized values were corrected for variation by normalizing against the mean of *cen-dg* and *cen-dh* as the *otr* is the region with the highest and most stable H3K9me2 enrichment (‘HC normalized’, *eq 2.1*). For ChIP experiments with anti-H3K36me3 and H3, input-normalized qPCR signals were normalized to the mean of 3 euchromatic loci (*act1*^+^, *ade2*^+^, *tef3*^+^) as an internal control, which was set to 1 (‘EC normalized’, *eq 2.2*). Using the mean of multiple euchromatic loci (‘EC’) instead of single locus (e.g. *act1*^+^) reduces bias coming from variations in ChIP experiments, especially when doing IP experiments with bulk histones.

**Transcript experimental normalization (RT-qPCR)**:

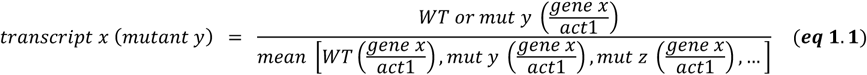

**Transcript mean WT normalization (RT-qPCR):**

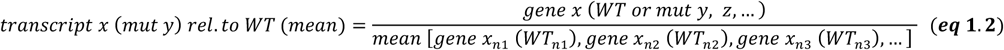

**Internal ChIP normalization H3K9me2 (ChIP-qPCR)**:

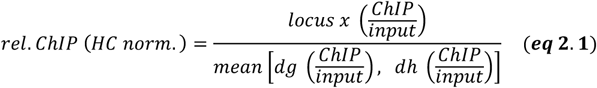

**Internal ChIP normalization H3K36me3 and H3 (ChIP-qPCR)**:

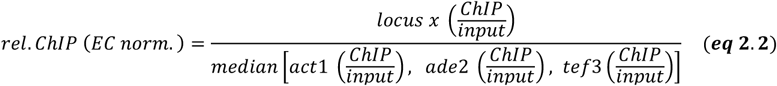

## Acknowledgements

We thank Marc Bühler for critical reading of the manuscript and the *brl1* point mutant fission yeast strains. This work was supported by grants awarded to SB from the German Research Foundation (BR 3511/2-1, BR 3511/4-1). SB is a member of the Collaborative Research Center 1064 funded by the German Research Foundation and acknowledges infrastructural support.

## Author contributions

PRG and SB designed the study. PRG generated yeast strains and performed RT-qPCR experiments with assistance by SFB. PRG and MC performed ChIP-qPCR experiments with assistance by SFB. MC performed silencing reporter assays. PRG and SB analyzed all data. SB wrote the manuscript, and PRG and MC contributed to editing.

---

**Supplementary Figure S1:**
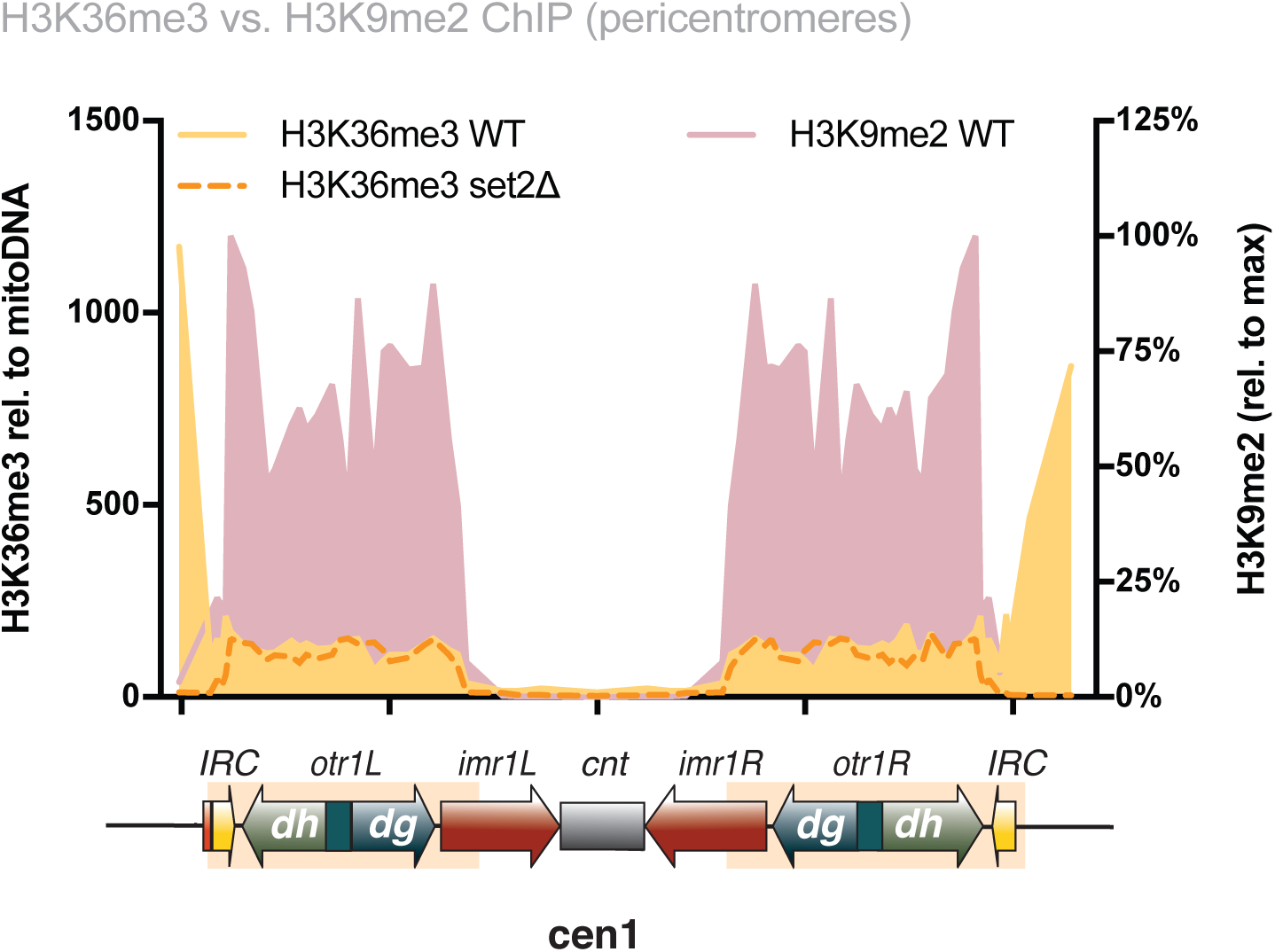
Cross reactivity analysis of anti-H3K36me3 to H3K9me2. Shown is a ChIP-qPCR analysis of a representative experiment using tiling primer arrays for H3K36me enrichments in WT and *set2*Δ cells at the pericentromeric region of chromosome 1. H3K36me3 enrichments were normalized to mitochondrial DNA, which is not affected by histone modifiers (left y-axis). For comparison with H3K9me2, a ChIP-qPCR analysis of WT cells (n=3) from Barrales et al. 2016 is shown (H3K9me enrichments normalized to maximal level in heterochromatin, which was set to 100%; right y-axis).

**Supplementary Table 1.** Yeast strains.

| STRAIN | SOURCE | IDENTIFIER |
| --- | --- | --- |
| *h<sup>+</sup> imr1L(NcoI)::ura4<sup>+</sup> otr1R(SphI)::ade6<sup>+</sup> leu1-32 ura4-DS/E ade6-M210* | (Braun et al., 2011) | PSB65 |
| *h<sup>+</sup> imr1L(NcoI)::ura4<sup>+</sup> otr1R(SphI)::ade6<sup>+</sup> leu1-32 ura4-DS/E ade6-M210 set2Δ::natMX* | This study | PSB1334 |
| *h<sup>-</sup> SPSQ (cyhR) SPL42 (cyhS) hphMX::cen1 imr1L(NcoI)::ura4<sup>+</sup> otr1R(SphI)::ade6<sup>+</sup> leu1-32 ura4-DS/E ade6-M210* | (Barrales et al., 2016) | PSB582 |
| *h<sup>-</sup> SPSQ (cyhR) SPL42 (cyhS) hphMX::cen1 imr1L(NcoI)::ura4<sup>+</sup> otr1R(SphI)::ade6<sup>+</sup> leu1-32 ura4-DS/E ade6-M210 pdp3Δ::natMX* | (Flury et al., 2017) | PSB689 |
| *h<sup>-</sup> SPSQ (cyhR) SPL42 (cyhS) hphMX::cen1 imr1L(NcoI)::ura4<sup>+</sup> otr1R(SphI)::ade6<sup>+</sup> leu1-32 ura4-DS/E ade6-M210 mst2Δ::natMX* | (Flury et al., 2017) | PSB1122 |
| *h<sup>-</sup> SPSQ (cyhR) SPL42 (cyhS) hphMX::cen1 imr1L(NcoI)::ura4<sup>+</sup> otr1R(SphI)::ade6<sup>+</sup> leu1-32 ura4-DS/E ade6-M210 nto1Δ::natMX* | This study | PSB1127 |
| *h<sup>-</sup> SPSQ (cyhR) SPL42 (cyhS) hphMX::cen1 imr1L(NcoI)::ura4<sup>+</sup> otr1R(SphI)::ade6<sup>+</sup> leu1-32 ura4-DS/E ade6-M210 ptf2Δ::natMX* | This study | PSB1130 |
| *h<sup>-</sup> SPSQ (cyhR) SPL42 (cyhS) hphMX::cen1 imr1L(NcoI)::ura4<sup>+</sup> otr1R(SphI)::ade6<sup>+</sup> leu1-32 ura4-DS/E ade6-M210 set2Δ::kanMX* | This study | PSB2111 |
| *h<sup>-</sup> SPSQ (cyhR) SPL42 (cyhS) hphMX::cen1 imr1L(NcoI)::ura4<sup>+</sup> otr1R(SphI)::ade6<sup>+</sup> leu1-32 ura4-DS/E ade6-M210 pdp3Δ::natMX set2Δ::kanMX* | This study | PSB2113 |
| *h<sup>-</sup> SPSQ (cyhR) SPL42 (cyhS) hphMX::cen1 imr1L(NcoI)::ura4<sup>+</sup> otr1R(SphI)::ade6<sup>+</sup> leu1-32 ura4-DS/E ade6-M210 nto1Δ::natMX set2Δ::kanMX* | This study | PSB2115 |
| *h<sup>-</sup> SPSQ (cyhR) SPL42 (cyhS) hphMX::cen1 imr1L(NcoI)::ura4<sup>+</sup> otr1R(SphI)::ade6<sup>+</sup> leu1-32 ura4-DS/E ade6-M210 mst2Δ::natMX set2Δ::kanMX* | This study | PSB2131 |
| *h<sup>-</sup> SPSQ (cyhR) SPL42 (cyhS) hphMX::cen1 imr1L(NcoI)::ura4<sup>+</sup> otr1R(SphI)::ade6<sup>+</sup> leu1-32 ura4-DS/E ade6-M210 ptf2Δ::natMX set2Δ::kanMX* | This study | PSB2325 |
| *h leu1-32 ura4-D18 ade6-704 trip1+::ade6<sup>+</sup> shade6-250/natMX* | (Flury et al., 2017) | PSB2356<br>(spb426) |
| *h<sup>-</sup> leu1-32 ura4-D18 ade6-704 trp1<sup>+</sup>::ade6<sup>+</sup> nmt1<sup>+</sup>::ade6-hp<sup>+</sup>::natMX brl1-K242R* | (Flury et al., 2017) | PSB2357<br>(spb2982) |
| *h leu1-32 ura4-D18 ade6-704 trip1+::ade6<sup>+</sup> shade6-250/natMX set2Δ::kanMX* | This study | PSB2361 |
| *h<sup>-</sup> leu1-32 ura4-D18 ade6-704 trp1<sup>+</sup>::ade6<sup>+</sup> nmt1<sup>+</sup>::ade6-hp<sup>+</sup>::natMX brl1-K242R set2Δ::kanMX* | This study | PSB2363 |

**Supplementary Table 2.**
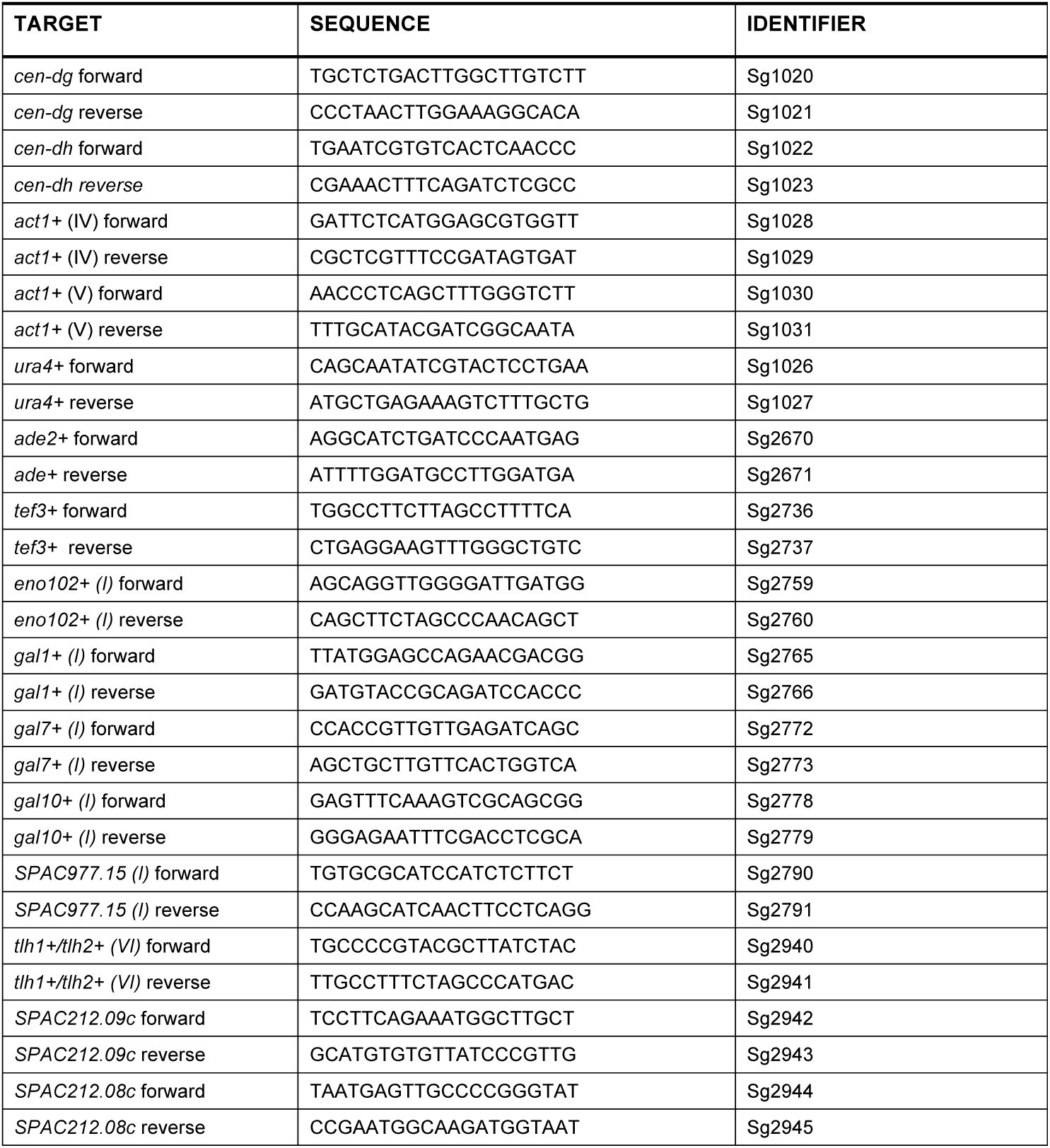
Oligonucleotides for Figures 1-4.

**Supplementary Table 3:**
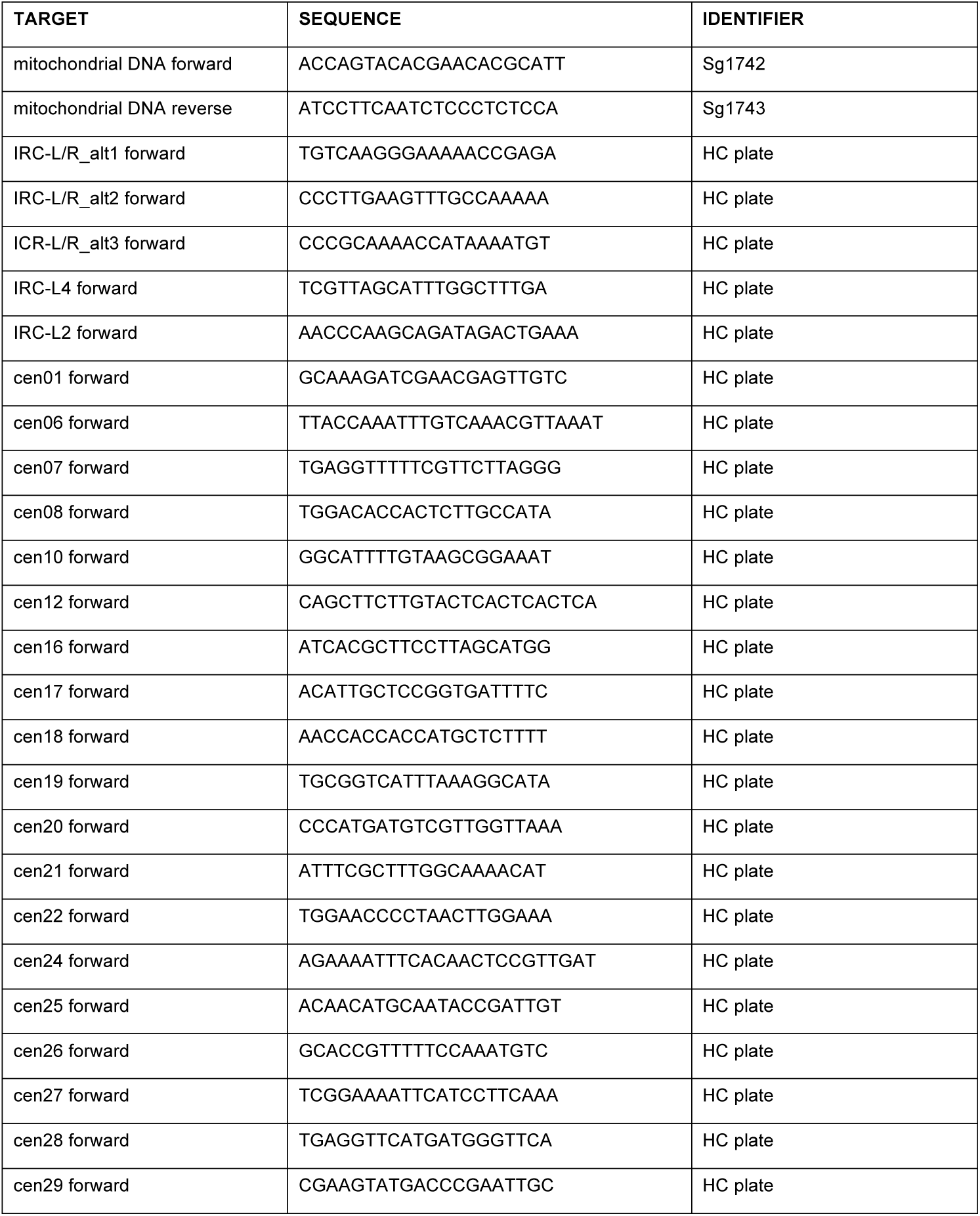
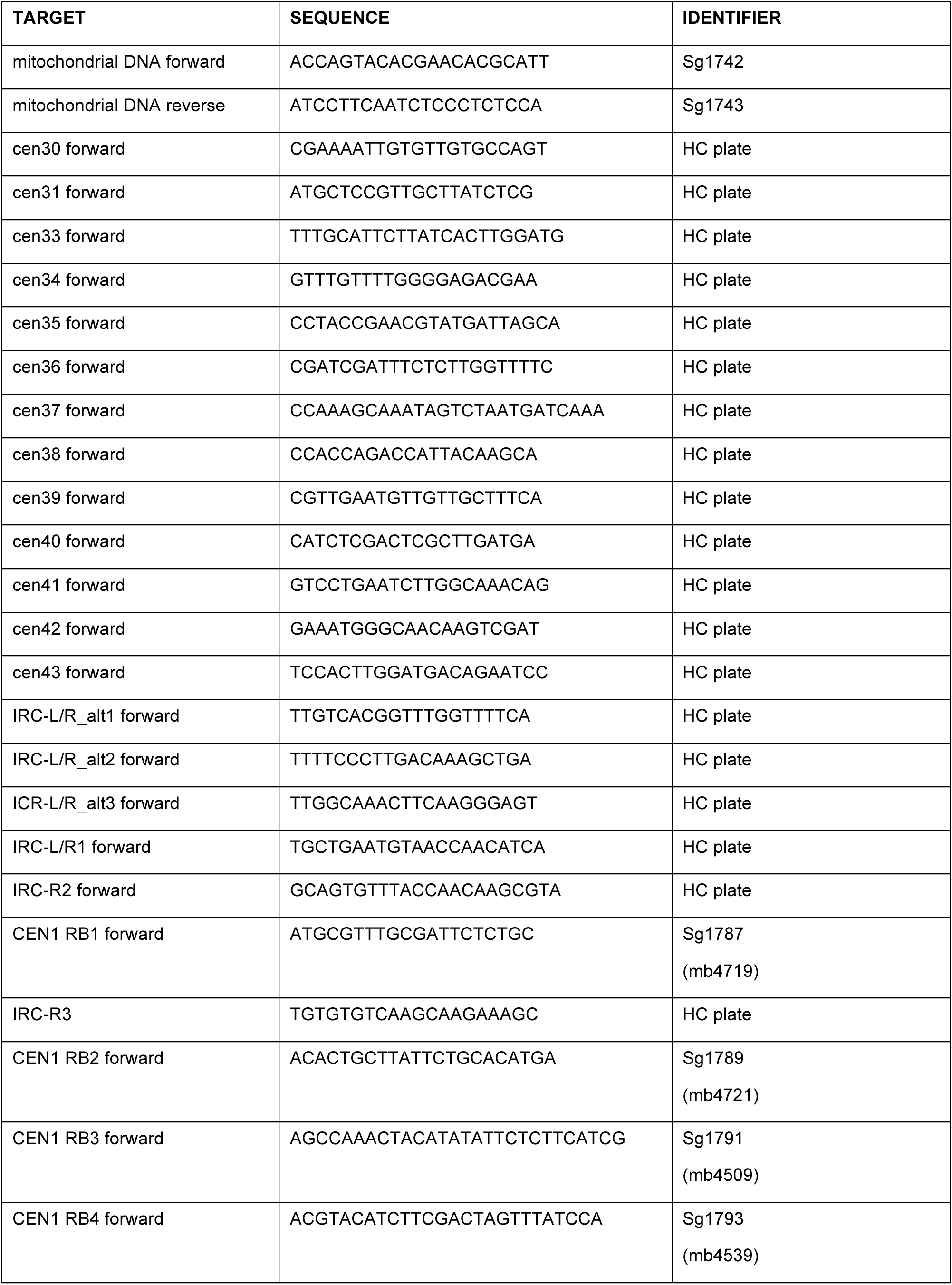
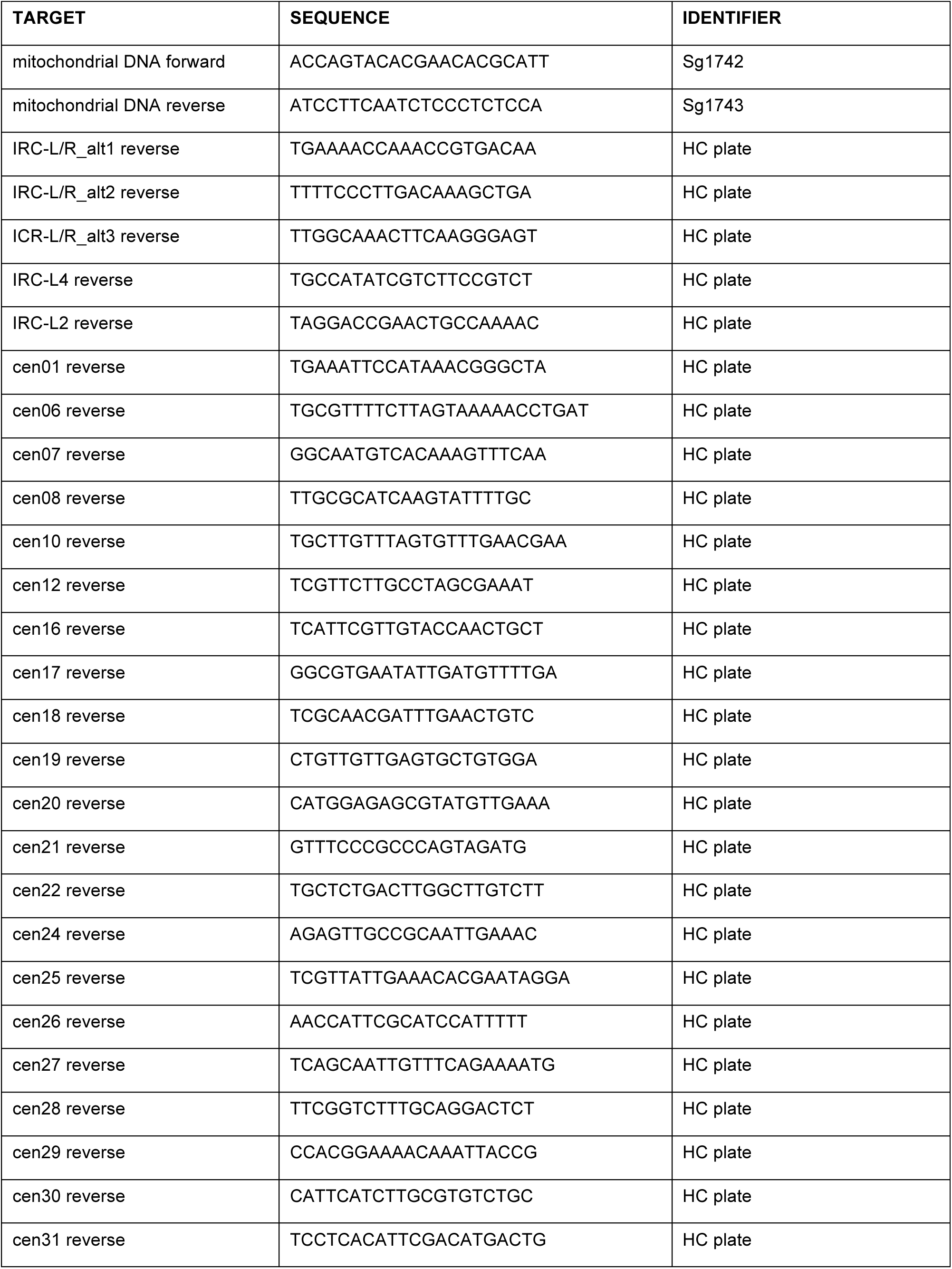
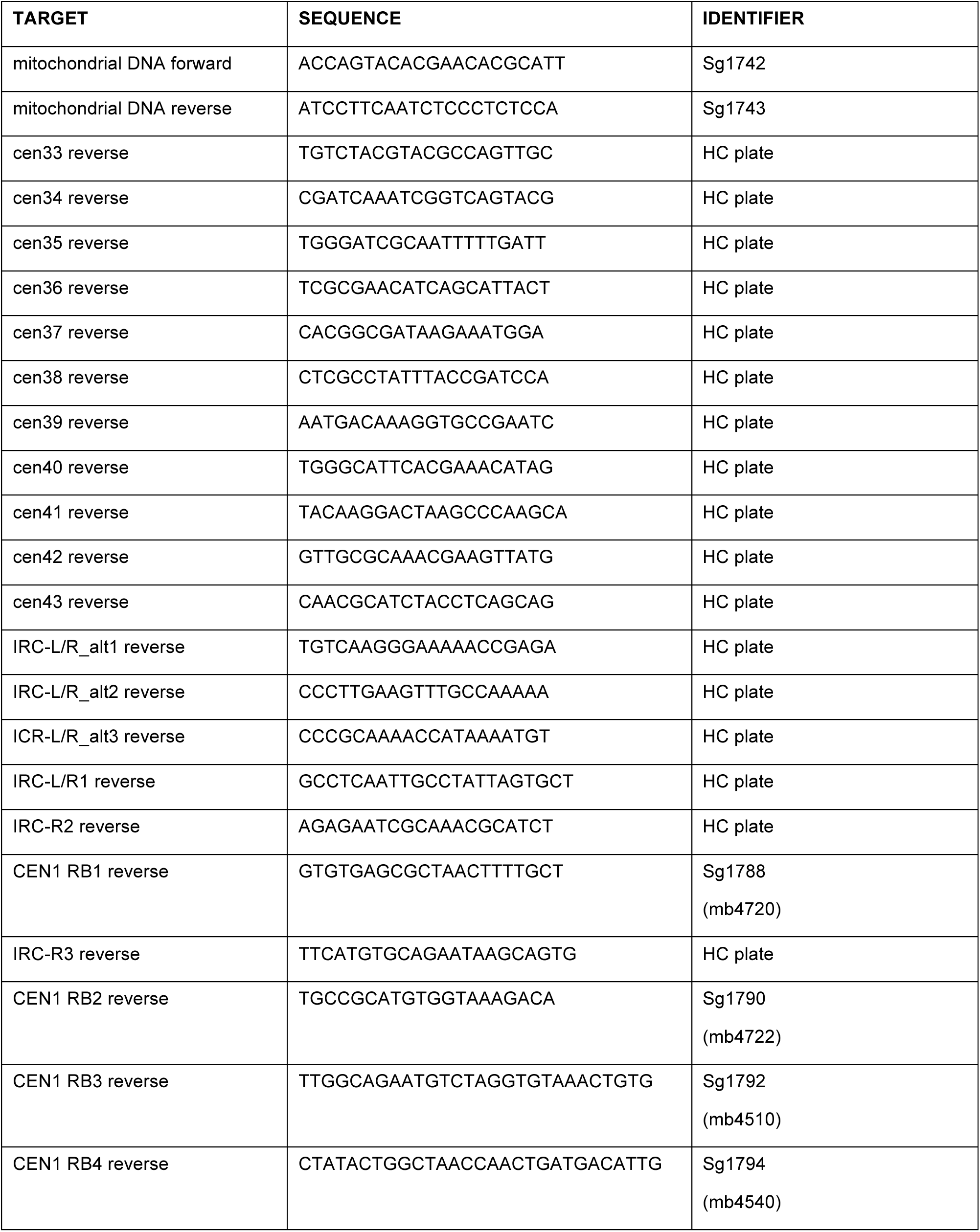
Oligonucleotides for Figure S1.

**Supplementary Table 4.** Antibodies.

| **ANTIBODY** | **SOURCE** | **IDENTIFIER** |
| --- | --- | --- |
| anti-H3K9me2 | Abcam | Cat# ab1220 |
| anti-H3K36me3 | Abcam | Cat# ab9050 |
| anti-H3 (ChIP) | Active Motif | Cat# 61475 |

**Supplementary Table 5.**
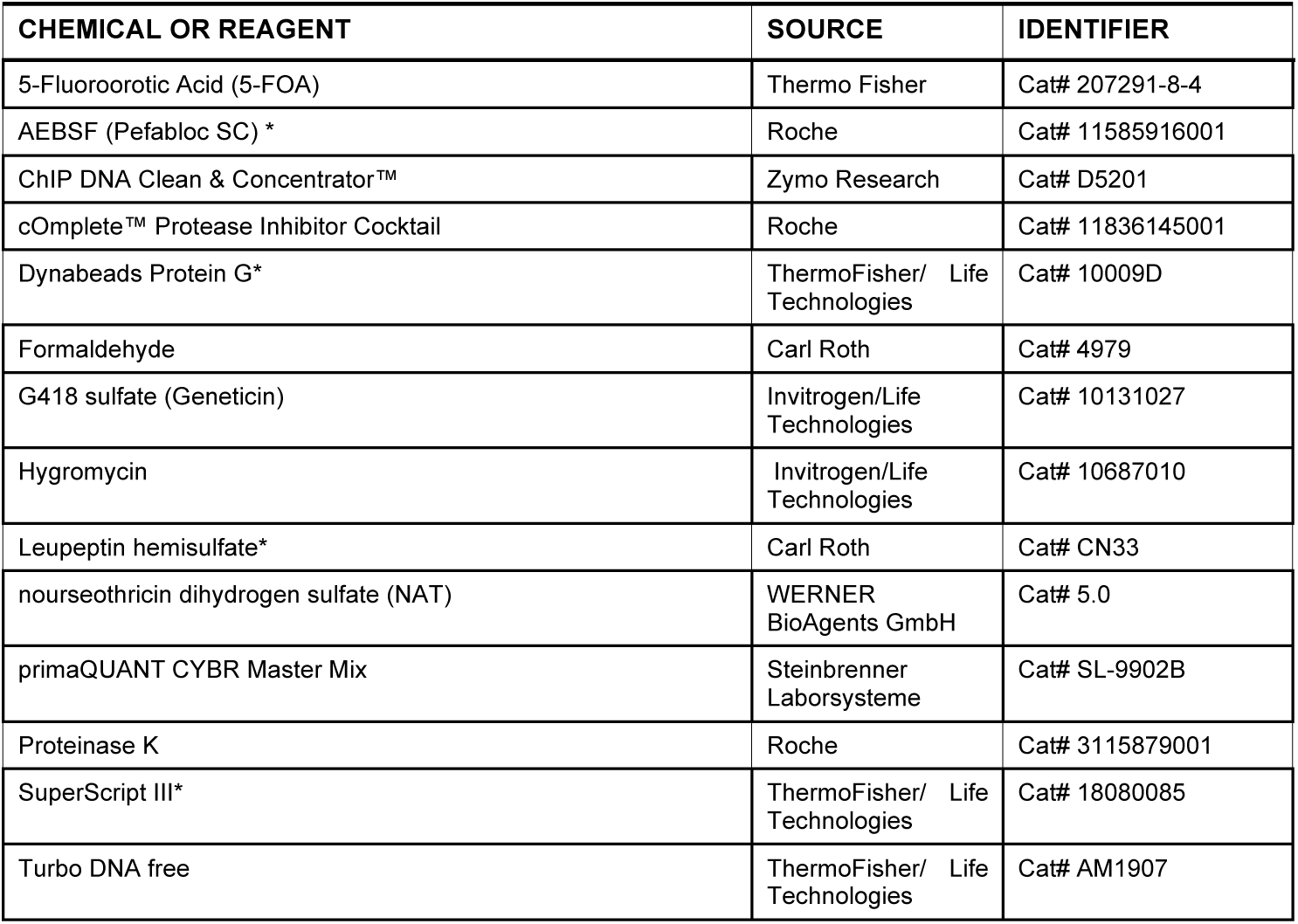
Chemicals, reagents and commercial assays.

| **CHEMICAL OR REAGENT** | **SOURCE** | **IDENTIFIER** |
| --- | --- | --- |
| 5-Fluoroorotic Acid (5-FOA) | Thermo Fisher | Cat# 207291-8-4 |
| AEBSF (Pefabloc SC) * | Roche | Cat# 11585916001 |
| ChIP DNA Clean & Concentrator™ | Zymo Research | Cat# D5201 |
| cOmplete™ Protease Inhibitor Cocktail | Roche | Cat# 11836145001 |
| Dynabeads Protein G* | ThermoFisher/ Life Technologies | Cat# 10009D |
| Formaldehyde | Carl Roth | Cat# 4979 |
| G418 sulfate (Geneticin) | Invitrogen/Life Technologies | Cat# 10131027 |
| Hygromycin | Invitrogen/Life Technologies | Cat# 10687010 |
| Leupeptin hemisulfate* | Carl Roth | Cat# CN33 |
| nourseothricin dihydrogen sulfate (NAT) | WERNER BioAgents GmbH | Cat# 5.0 |
| primaQUANT CYBR Master Mix | Steinbrenner Laborsysteme | Cat# SL-9902B |
| Proteinase K | Roche | Cat# 3115879001 |
| SuperScript III* | ThermoFisher/ Life Technologies | Cat# 18080085 |
| Turbo DNA free | ThermoFisher/ Life Technologies | Cat# AM1907 |

